# Single cell resolution analysis of multi-tissue derived human iNKT cells reveals novel transcriptional paradigms

**DOI:** 10.1101/2024.03.22.583992

**Authors:** Reyka G. Jayasinghe, Derek Hollingsworth, Chaiyaporn Boonchalermvichian, Biki Gupta, Hao Yan, Jeanette Baker, Beruh Dejene, Kenneth I Weinberg, Robert S. Negrin, Melissa Mavers

## Abstract

Invariant natural killer T (iNKT) cells are evolutionarily conserved innate lymphocytes important for host defense against pathogens. Further, they are increasingly recognized to play a role in tumor immune surveillance and in protection against graft versus host disease, and they are of particular importance as a universal donor for cellular therapies. Therefore, a thorough understanding of the biology of iNKT cells is critical. Murine studies have revealed the existence of transcriptionally and functionally distinct subsets, similar to T helper cell subsets. However, a comprehensive study of human iNKT cell heterogeneity is lacking. Herein, we define the transcriptomic heterogeneity of human iNKT cells derived from multiple immunologically relevant tissues, including peripheral blood, cord blood, bone marrow, and thymus, using single cell RNA-sequencing. We describe human iNKT cells with a naïve/precursor transcriptional pattern, a Th2-like signature, and Th1/17/NK-like gene expression. This combined Th1/17 pattern of gene expression differs from previously described murine iNKT subsets in which Th1- and Th17- like iNKT cells are distinct populations. We also describe transcription factors regulating human iNKT cells with distinct gene expression patterns not previously described in mice. Further, we demonstrate a novel T effector memory RA^+^ (TEMRA)-like pattern of expression in some human iNKT cells. Additionally, we provide an in-depth transcriptional analysis of human CD8^+^ iNKT cells, revealing cells with two distinct expression patterns—one consistent with naïve/precursor cells and one consistent with Th1/17/NK-like cells. Collectively, our data provide critical insights into the transcriptional heterogeneity of human iNKT cells, providing a platform to facilitate future functional studies and to inform the development of iNKT-based cellular therapies.

## INTRODUCTION

Invariant natural killer T (iNKT) cells lie at the interface of innate and adaptive immunity and play important protective roles in immune responses to pathogens and surveillance for malignant cells. iNKT cells recognize glycolipid antigen via their invariant T cell receptor, comprised of Vα24Jα18 paired with Vβ11 in humans and Vα14Jα18 paired with Vβ2, 7, or 8 in mice^1^. This recognition requires presentation of the glycolipid antigen by the major histocompatibility complex (MHC)-like protein CD1d^2^. Because CD1d is monomorphic, iNKT cells are incapable of exerting pathogenic non-self MHC recognition and are therefore unable to initiate graft-versus-host disease (GVHD). This lack of allorecognition makes iNKT cells a powerful platform for universal donor cellular therapies, with many advantages over conventional T cells, and a clinical trial of allogeneic chimeric antigen receptor (CAR)-engineered iNKT cells for cancer therapy is underway (NCT03774654).

While iNKT cells are not capable of causing GVHD, substantial evidence suggests that they can suppress it^3^. The adoptive transfer of even very low numbers of donor or third party iNKT cells into mouse models of acute and chronic GVHD provides protection from disease^4,5^. The use of conditioning regimens which facilitate the relative retention of host iNKT cells is also associated with protection from GVHD in both mice and humans^6–9^. Clinical studies in hematopoietic stem cell transplantation (HSCT) have demonstrated that donors who have higher iNKT cell numbers in the graft^10^, or more rapid reconstitution of iNKT cells post-transplant, are associated with reduced GVHD as well.

Importantly, however, functionally distinct subsets of iNKT cells have been reported in mice. These subsets have been described as similar to T helper cell subsets, with Th1-like iNKT1 cells expressing T-bet and producing primarily interferon (IFN)γ, Th2-like iNKT2 cells expressing the highest levels of promyelocytic leukemia zinc finger (PLZF) (albeit with lower levels of expression variably retained in other subsets) and primarily producing interleukin (IL)-4, and Th17-like iNKT17 cells expressing RAR-related orphan receptor-gamma (RORγt) and producing primarily IL-17^1,11–15^. iNKT2 and iNKT17 cells were shown to suppress acute GVHD in murine models, while iNKT1 cells demonstrated the most cytotoxic function toward malignant cells^12^. These results suggest that the safety and efficacy of iNKT cell-based cellular therapies might be enhanced by aligning the use of particular subsets with the goal of the cellular therapy. For example, CAR-iNKT products could be developed from purified iNKT1 cells, while other subsets might be better suited for immunosuppressive cellular therapies (ie. to prevent GVHD). However, a better understanding of human iNKT subsets is required to realize the full clinical potential of iNKT cells, and a comprehensive study of human iNKT heterogeneity with robust cell numbers and tissue sources is lacking.

Herein, we define the transcriptomic heterogeneity of human iNKT cells derived from multiple immunologically relevant tissues, including peripheral blood, cord blood, bone marrow, and thymus. Because future therapeutic use of distinct subsets would require the ability to distinguish these subsets in an unstimulated state, we specifically sought to study the heterogeneity of unstimulated human iNKT cells. While identifying many of the same populations described previously in mice, we report several distinct differences in transcriptomic patterns of human iNKT cells. These novel findings include a mixed Th1/17 signature, two transcriptionally distinct CD8^+^ iNKT cell clusters, and a CD4^-^CD8^-^ T effector memory RA^+^ (TEMRA)-like subcluster. These results define the transcriptional heterogeneity of human iNKT cells and provide a critical foundation to build our understanding of human iNKT cell subsets.

## RESULTS

### Surveying the landscape of human iNKT cells at single-cell resolution

The transcriptional heterogeneity of human iNKT cells was investigated in two experiments, one with 11 unpaired donor samples of peripheral blood (PB, n=3), cord blood (CB, n=3), thymus (n=3), and bone marrow (BM, n=2) and another with PB only (n=10) (Fig 1A-B). iNKT cells were enriched by magnetic beads and fluorescence activated cell sorted (FACS) using antibodies to the invariant T cell receptor. CD24^+^ stage 0 iNKT precursor cells were removed via FACS to ensure adequate numbers of differentiating and mature thymic iNKT cells to better facilitate comparison across tissues. In total, 13,047 cells passed quality control for single cell RNA-sequencing (scRNA-seq) in the multi-tissue experiment. A small cluster (representing <2% of the total) significantly upregulated *FCER1G, FCGR3A, LYN,* and *IL1B* identifying the cells as macrophages and was excluded from further analysis. A total of 24,758 cells passed quality control for scRNA-seq in the peripheral blood only experiment. The multi-tissue experiment utilized whole transcriptome analysis (WTA) as well as targeted scRNA-seq using a 435 gene panel custom designed to include genes important for natural killer (NK) cell and T cell function, including both proinflammatory and immune regulatory subsets (Supp Table 1). Cells in the peripheral blood only experiment underwent targeted sequencing only. We also utilized oligomer-conjugated antibodies to detect CD45RA, CD45RO, CD4, and CD8.

**Figure 1.**
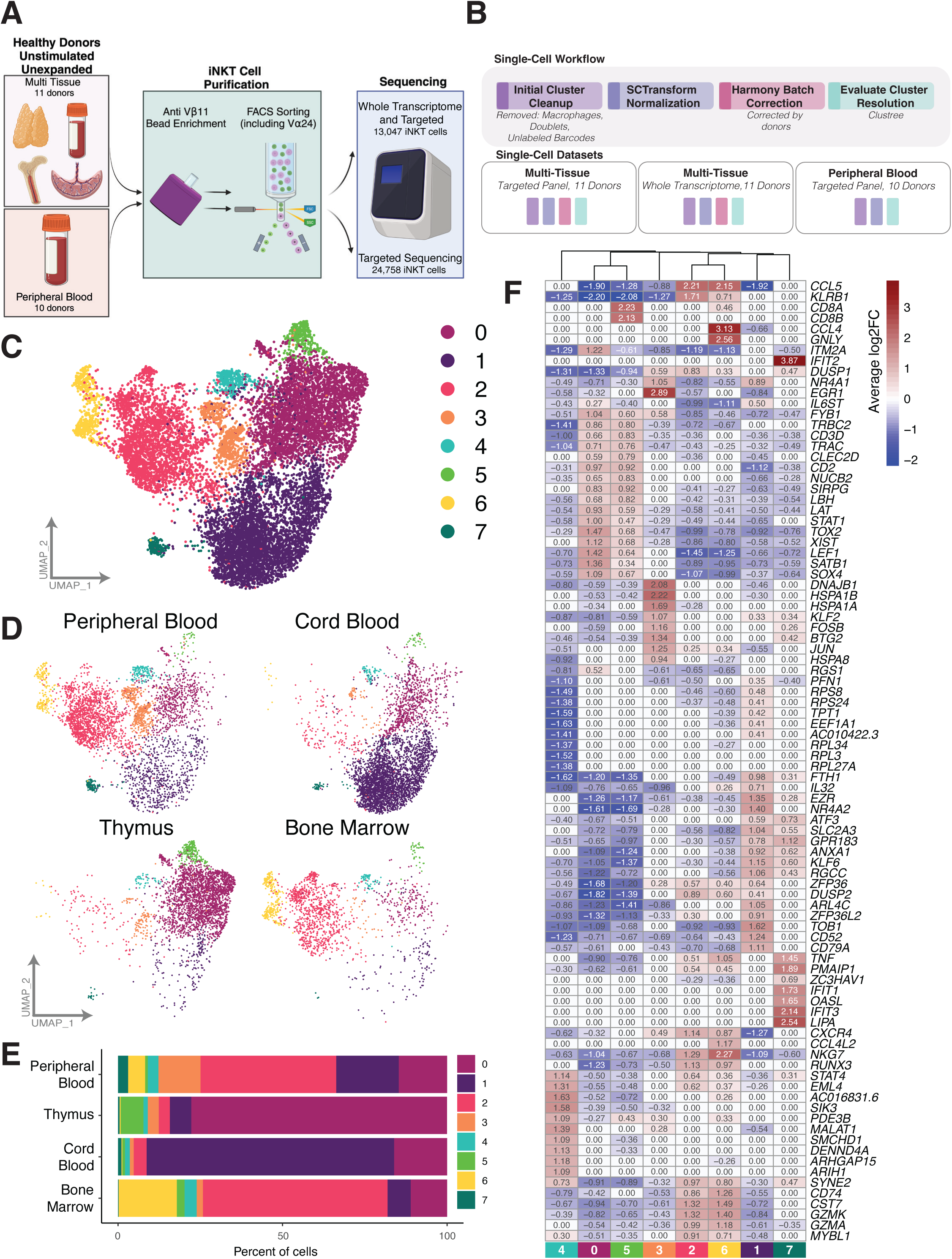
Overview of scRNA-seq datasets generated from human iNKT cells. A) Schematic of iNKT cell purification and sequencing. B) Overview of single-cell workflow for each dataset. C) UMAP of all iNKT cells isolated from the multi-tissue cohort. Cells are colored by cluster assignment. D) UMAP representation of all iNKT cells separated by tissue source. Cells are colored by cluster assignment. E) Barplot indicating proportions of iNKT cells restricted to each cluster separated by tissue source. F) Heatmap of the top 10 differentially expressed genes identified for each cluster. Each cell is colored by the average log2FC of each gene identified in each cluster. A value of 0 indicates the gene was not identified as being significantly up or down regulated for the labeled cluster.

### Patterns of *TBX21, ZBTB16,* and *RORC* expression in human iNKT cells reveal Th1/17-like cells not described in mice

Transcriptional heterogeneity of human iNKT cells from PB, CB, thymus, and BM revealed eight iNKT cell clusters in unsupervised clustering analysis (Fig 1C). While iNKT cells from some tissue sources dominated certain clusters, peripheral blood-derived iNKT cells were relatively equally distributed among all clusters (Fig 1D-E). These findings persisted across individual donors (Supp Fig 1A). For each cluster, the top 10 defining markers were visualized, revealing distinguishing features between clusters (Fig 1F, Supp Table 2).

Because murine iNKT cells can be clearly divided into functional subsets based on their transcription factor expression patterns, with iNKT1 cells exclusively expressing *Tbx21*, iNKT17 cells exclusively expressing *Rorc*, and iNKT2 cells lacking *Tbx21* and *Rorc* and expressing the highest levels of *Zbtb16*^1,11–13^, we next sought to evaluate the expression patterns of these transcription factors in human iNKT cells. In contrast to murine cells, we found that clusters of human iNKT cells which express *TBX21*, Cluster 2 (C2) and C6, display co-expression of *RORC*, representing a Th1/17-like signature (Fig 2A). These clusters were predominated by BM-derived iNKT cells (Fig 1D-E, 2A). C1 had upregulation of *ZBTB16* with downregulation of *TBX21* and *RORC*, consistent with a Th2-like iNKT cell signature. C4 and C7 co-expressed *ZBTB16* and *TBX21*, though *TBX21* was downregulated compared to C2 and C6, suggesting these clusters may represent transitional cells between Th2-like iNKT cells and Th1/17-like iNKT cells (Fig 2A). This correlates with previous work in murine iNKT cells demonstrating that iNKT1 cells differentiate from iNKT2 cells, with transitional populations in between^16^. CB-derived iNKT cells predominated in those clusters with highest *ZBTB16* expression (Fig 1D-E, Fig 2A). Three clusters, C0, C3, and C5, had minimal expression of *TBX21* and *RORC* and retained expression of *ZBTB16*, albeit relatively downregulated relative to clusters C1 and C4 (Fig 2A). These cells were largely thymic-derived and may represent early iNKT cells differentiating towards Th1/17-like iNKT cells, correlating with data from a recent murine study^17^ (Fig 1D-E). Collectively, these results suggest that human iNKT cells can be described as Th1/17-like, Th2-like, transitional, or early differentiating cells based on their expression patterns of transcription factors canonically associated with murine iNKT cells.

**Figure 2.**
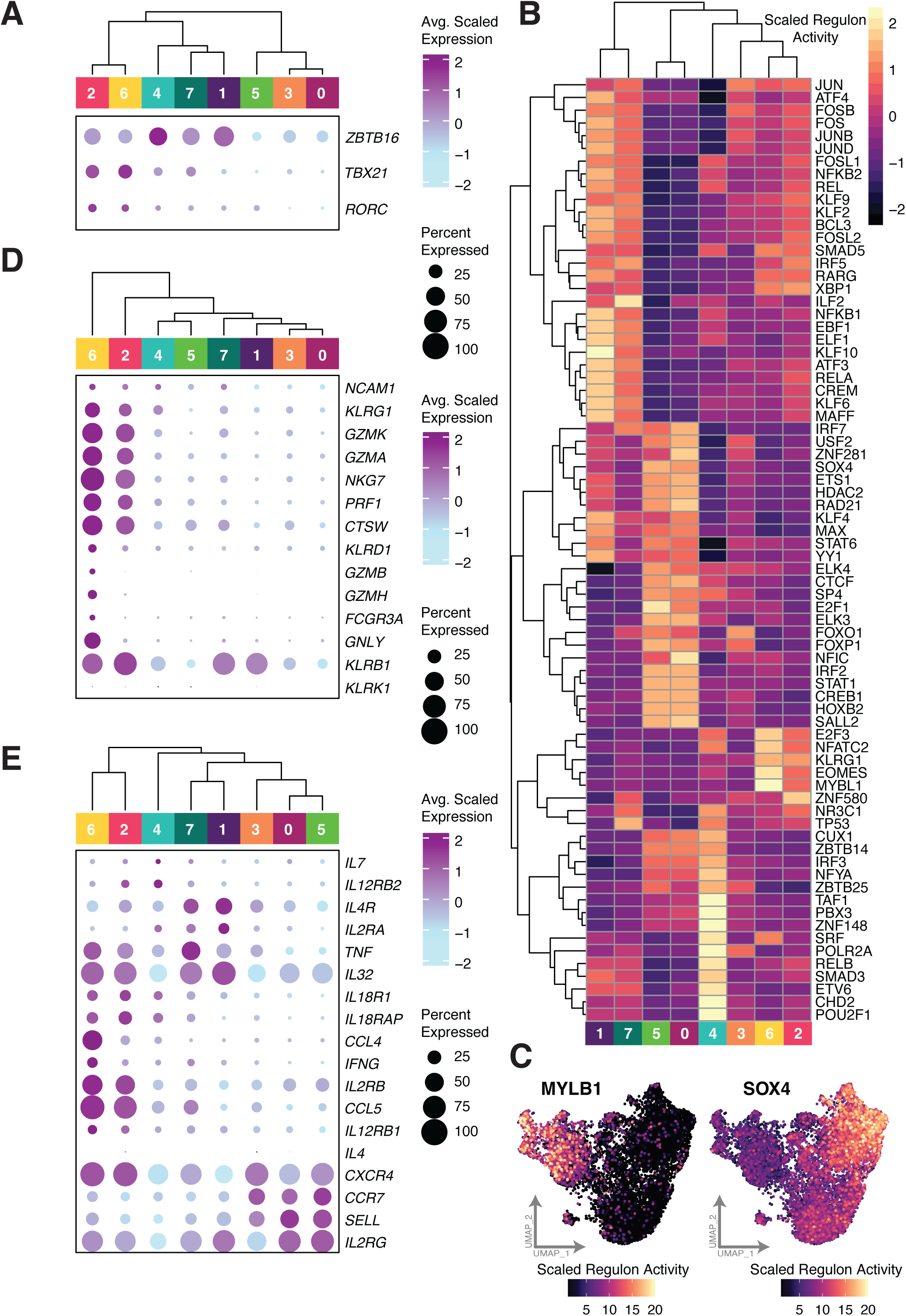
Human iNKT cell transcriptional heterogeneity reveals Th1/17/NK-like cells, Th2-like cells, and early differentiating cells. A) Expression dot plots of key transcription factors. Size of dot indicates percent of cells expressing the indicated gene and color indicates the average scaled expression. B) Heatmap of differentially expressed regulons for each iNKT cluster. Each cell is colored by the scaled regulon activity of all cells belonging to the annotated cluster. C) UMAPs of two key regulons identified. Each cell is colored by the scaled regulon activity score of the indicated regulon. D-E) Expression dotplot of a subset of genes. Size of dot indicates percent of cells expressing each gene and color indicates the average scaled expression.

### Regulon analysis reveals transcription factors that may play a critical role in distinct human iNKT subset functions

We next performed a regulon analysis to unveil additional transcription factors regulating distinct clusters of human iNKT cells. Regulon analyses measure the change in transcript expression of all targets of a given transcription factor to understand the activity of that transcription factor in the cell. By evaluating regulons that are differentially expressed by cluster using SCENIC, we identified several transcription factors with upregulated regulons in each cluster (Fig 2B, Supp Table 3). Regulons with the largest average log2 fold change between the indicated cluster and all other clusters included MYBL1 and EOMES in C2 and C6, EBF1, JUND and ATF3 regulons in C1, and SOX4 and FOXO1 regulons in C0 and C5 (Fig 2B-C). The EOMES regulon contains 33 target genes including *GZMK*, *NKG7*, *KLRG1*, *CCL4* and *LAG3* all of which are upregulated in C2 and C6. Eomesodermin (protein encoded by *Eomes*) has been shown to regulate iNKT cell differentiation into iNKT1 cells and KLRG1-expressing iNKT cells in mice^18^. *MYBL1* has been associated with innateness in lymphocytes^19^, and the MYBL1 regulon contains the target gene *NKG7* which is upregulated in iNKT1 cells in mice^16,20^. Notably, the *MYBL1* gene itself is differentially upregulated in C2 and C6 in our analysis (Fig 1F, Supp Table 2). *MYBL1* is also contained within the regulon of FOSL2. The FOSL2 regulon is upregulated in C1 and C7, suggesting a sequential activation of transcription factors as cells differentiate from Th2-like into Th1-like through the transitional C7. Several regulons (including E2F3 and NFATC2) are shared between C4 and C2/C6, further highlighting the transitional role of C4. *EBF1* expression is associated with B cell development^21^ and, interestingly, the EBF1 regulon is differentially upregulated in C1 which also exhibits high expression of CD79A (C1: 1.11 avg log2FC). CD79A is vital for B cell development and surface expression of the B cell receptor^22,23^ and is also transcriptionally upregulated in a subset of human peripheral blood iNKT cells^24^. The CD79A gene is contained within the regulons of the following transcription factors: ATF3, CREM, FOS, FOSB, FOSL2 and JUND; all of which contain high SCENIC activity scores in C1. Analysis of differentially expressed genes using the Ingenuity Pathway Analysis (IPA) Tool revealed C1 has activation of Signaling by the B Cell Receptor (BCR) pathway (z.score= 2.31, Supp Fig 2A, Supp Table 4). The specific role of these B cell developmental factors in iNKT cells remains to be elucidated. Another regulon upregulated in C1 is ATF3 which negatively regulates IFNγ expression in murine NK cells^25^, further suggesting a Th2-skewing of this cluster. IPA analysis further supports the Th2 Pathway is activated in C1 (z.score= 1.21, Supp Fig 2A, Supp Table 4). Interestingly, TBX21 was not identified in our regulon analysis. While the ZBTB16 regulon also does not reach significance in our analysis, this gene is contained within the target list of 13 other differentially regulated transcription factors that are upregulated in activity in C1 and C7 (Fig 2B, Supp Table 3). Collectively, these transcriptomic data reveal some similarities in murine and human iNKT cell transcription factor gene expression, while highlighting novel transcription factors important in driving human iNKT cell transcriptional diversity.

### Molecular expression signatures suggest distinct effector functions between iNKT cell clusters

Earlier research in murine studies has indicated that iNKT1 cells possess cytotoxic capabilities similar to cytotoxic T and NK cells and absent in other iNKT subsets. To further investigate this in human iNKT cells, we interrogated the expression pattern of genes important for cytotoxicity and other NK cell functions. Th1/17-like C2 and C6 have significantly upregulated expression of many of these genes relative to other clusters (Fig 2D), suggesting that these clusters are characterized by cytotoxic function. This included high expression of *GZMK* (gene encoding granzyme K) in most cells, with a very small proportion of cells expressing *GZMB* in C6 only (*GZMK* - C2:1.32 avg log2FC C6: 1.40 avg log2FC), similar to what has been reported in many cytotoxic innate lymphocytes^26^. We also observed increased expression of *KLRG1* and *GZMA* in these two clusters, similar to a KLRG1^+^GZMA^+^ population of long-lived iNKT cells deriving from iNKT1 cells in mice (*KLRG1* – C2: 0.47 C6: 0.57 avg log2FC; *GZMA* – C2:0.99 C6:1.18 avg log2FC)^27^. Expression of *GNLY* (gene encoding granulysin) was strongly upregulated in C6 relative to other clusters (2.56 avg log2FC Fig 1F). Granulysin is a cytolytic component of cytotoxic T and NK cell granules which also possesses chemoattractant function and induces expression of a number of inflammatory cytokines^28^. *KLRB1* (gene encoding CD161, the human homolog of NK1.1 in C57BL/6 mice) has been described as a marker of fully differentiated iNKT cells corresponding to iNKT1 cells^29,30^. Accordingly, *KLRB1* expression arises in C1 and C7 and maintains the highest expression in C2 and C6 (C2: 1.71; C6: 0.71 avg log2FC). Pathway analysis revealed increased Natural Killer Signaling and Interferon Gamma Signaling in C2 and C6 relative to other clusters (Supp Fig 2A, Supp Table 4). Overall, these results suggest that the Th1/17-like clusters (C1 and C6) of unstimulated human iNKT cells can be best described as Th1/17/NK-like. Despite this, we found a very low percentage of cells expressing *NCAM1* (<7%, gene encoding CD56) across all clusters, though within that low percentage, gene expression was highest in C2, C4, and C6 as compared to other clusters (Fig 2D). A previous report measuring CD56 at the protein level similarly found that very few iNKT cells defined by expression of the invariant TCR express CD56^31^.

Detection of cytokines is notoriously difficult in scRNA-seq analyses, particularly in unstimulated cells, and both *IL4* and *IFNG* were sparsely detected, though *IFNG* was most significantly upregulated in C6 (Fig 2E, Supp Fig 2B). *TNF* expression was highest in C2, C6, and C7. Cytokine receptor expression suggested functionally distinct profiles of cytokine responsiveness between clusters, including relatively increased expression of *IL18R1*, *IL18RAP, IL12RB1, and IL12RB2* in C2 and C6, suggesting further similarity to NK cells (Fig 2E)^32^. Expression of *IL2RB* (gene encoding IL2Rβ, part of the intermediate and high affinity IL-2 receptors as well as the IL-15 receptor) was also upregulated in these two clusters (C2: 0.90 avg log2FC, C6: 0.87 avg log2FC), although *IL15RA* was undetectable in our analysis. We also noted *IL4R* expression was highest in C1 (0.40 avg log2FC), further reflecting the Th2-skewed signature of these cells, and slightly decreased in C7 relative to C1, denoting the transitional nature of this cluster. This correlates with previous studies demonstrating that cord blood and neonatal derived iNKT cells have a Th2-skewed cytokine production pattern^33,34^. C7 was also characterized by increased *IFIT1*, *IFIT2*, and *IFIT3* expression (Fig 1F), indicative of Type I IFN signaling important for innate immune responses^35^. Pathway analysis reveals IFNα/β signaling is highly enriched in C1 and, to a lesser extent, in C7, suggesting the sequential activation of IFNα/β signaling leading to expression of *IFIT* targets and further supporting the transitional profile of C7 (z.score = 2.44, Supp Fig 2A, Supp Table 4). *CCL5* and *CXCR4* were the most highly expressed chemokine and chemokine receptor, respectively, and were upregulated in C2 and C6 (*CCL5*: C2: 2.20, C6: 2.15 avg log2FC; *CXCR4*: C2:1.13 C6:0.86 avg log2FC) (Fig 2E, Supp Fig 2B). Chemokine Signaling pathways were also activated in C2 (z.score = 0.71) and C6 (z.score = 1.41) in the Ingenuity Pathway analysis (Supp Fig 2A). CXCR4 plays a chemotactic role for lymphocytes via its role as a receptor for CXCL12/stromal-derived factor 1 and was previously shown to be highly expressed in peripheral blood iNKT cells^36^. CCL5 recruits many immune cells to sites of inflammation and is upregulated in a subset of iNKT1 cells in mice^16^. Collectively, these results provide additional support for the Th1/17/NK-like signature of C2 and C6 and the Th2-like signature of C1.

### Expression patterns reveal unique T effector memory CD45RA^+^ (TEMRA)-like cells and novel description of CD8^+^ human iNKT cells at the transcriptional level

The inclusion of oligomer-conjugated antibodies allowed for specific detection of CD45RA and CD45RO in our scRNA-seq analysis. Together with analysis of mRNA expression of *CCR7* and *SELL* (gene encoding CD62L), this allowed for the evaluation of naïve and memory phenotypes of human iNKT cells, following the canonical definition of these populations in human conventional T cells. The early differentiating clusters (C0, C3, and C5) had upregulation of *CCR7* and *SELL*, with relatively increased CD45RA, similar to naïve T cells, suggesting a naïve/precursor signature in these clusters (Fig 3A). This is further supported by the relative upregulation of genes known to play important roles in iNKT cell development, including *LEF1, TOX2, SATB1, SOX4,* and *ITM2A* (Fig 1F)^16,20,37–40^. Indeed, previous reports showed that lymphoid enhancer binding factor 1 (LEF1) expression correlated with, and appeared to regulate, CD62L expression in *ex vivo* expanded iNKT cells, albeit with a central memory phenotype possibly reflecting changes occurring during culture^41,42^. The majority of the rest of the iNKT cells in our analysis displayed an effector memory signature with no/low expression of *CCR7* and CD45RA. Surprisingly, however, we detected that a subset of cells within C6 (designated subcluster 6.2, C6.2) predominantly expressed CD45RA with absent *CCR7* (and *SELL*) transcript, similar to that of TEMRA cells, which in conventional T cells have been described as memory cells with re-expression of CD45RA (Fig 3A)^43^.

**Figure 3.**
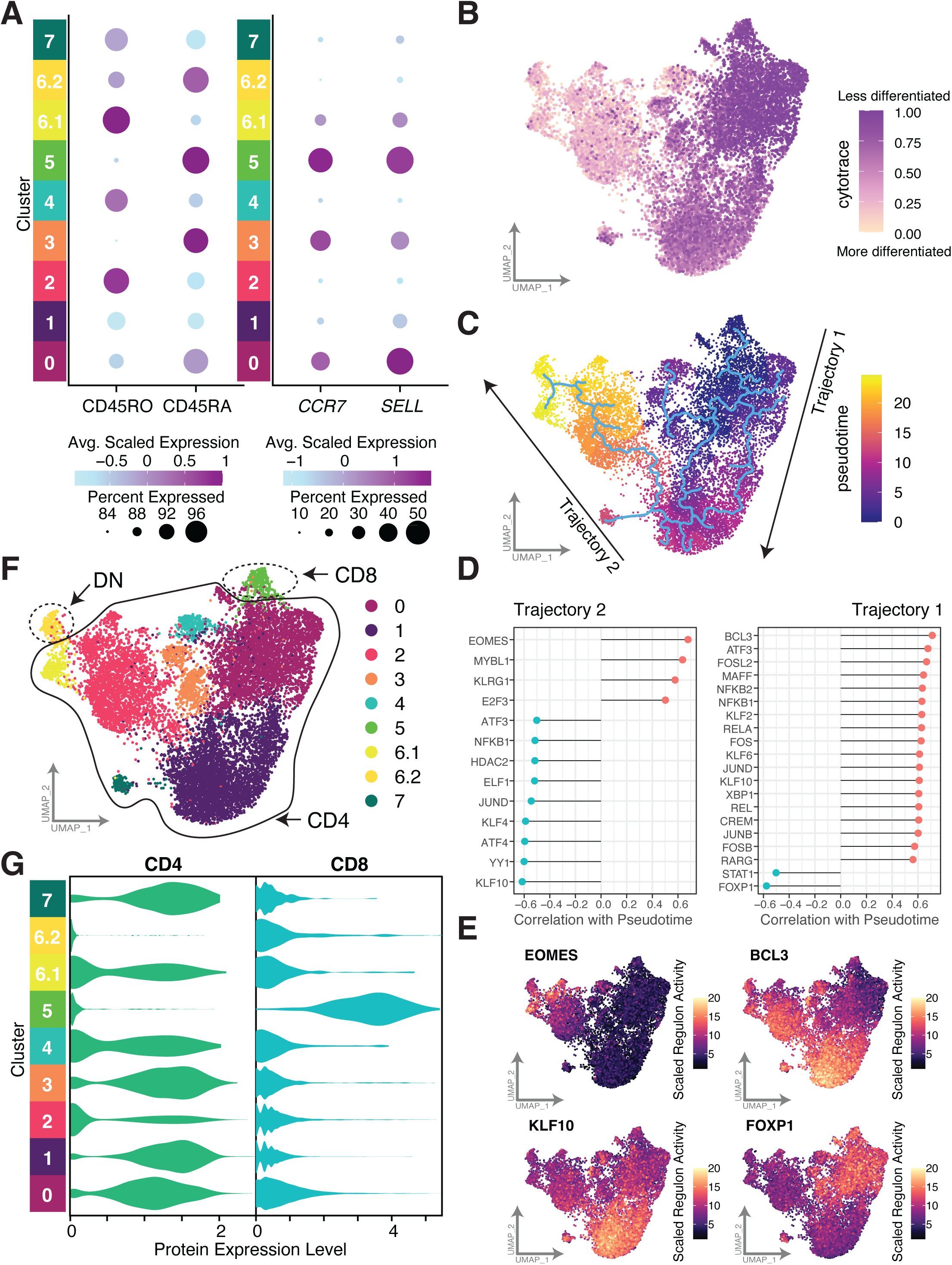
Naïve/memory expression patterns and differentiation trajectories demonstrate novel T effector memory CD45RA^+^ (TEMRA)-like iNKT cells. A) Dot plot of CD45RO and CD45RA protein expression and *CCR7* and *SELL* gene expression. For each figure, size of dot indicates percent of cells expressing the protein/gene of interest and color indicates average scaled expression. B) UMAP of iNKT cells colored by CytoTRACE score. C) UMAP of iNKT cells colored by pseudotime. Overlaying trace indicates predicted trajectories using Monocle3. Two arrows indicate annotated trajectories that will be highlighted separately in the following figures. D) Regulons correlating with pseudotime across the annotated trajectory if > 0.5 or < -0.5. For each gene, the resulting correlation value (after FDR correction) of the regulon with pseudotime is indicated. E) UMAP representation of annotated regulon with each cell colored by SCENIC activity score of the top correlated and anti-correlated gene from trajectory1 and trajectory2. F) UMAP of iNKT cells colored by cluster assignment. Regions of UMAP are circled based on observed protein expression of CD4 and CD8. G) Violin plot of CD4 and CD8 protein expression separated by cluster.

To further evaluate the differentiation trajectory of human iNKT cells, we performed Cellular Trajectory Reconstruction Analysis using gene Counts and Expression (CytoTRACE) as well as pseudotime trajectory analysis (Monocle3) (Fig 3B-C). CytoTRACE predicts the differentiation state of cells from an scRNA-seq-based determinant of developmental potential, assuming that transcriptional diversity (the number of different genes expressed in a cell) decreases during differentiation. In a complementary fashion, pseudotime infers the order of cells along a trajectory or related lineage. CytoTRACE analysis revealed that C0 represented the least differentiated cluster, correlating with our naïve/memory signature data and transcription factor expression data, and we set this as the root for pseudotime. Thymic derived cells contributed the largest number of cells to the naïve/precursor cluster (C0) providing further evidence to the starting point for pseudotime analysis validated with the CytoTRACE analysis. Differentiation also appeared to progress through the Th2-like cluster (C1) to the Th1/17/NK-like clusters (C2 & C6).

Next we evaluated regulons that correlated with the differentiation state (pseudotime) from naïve/precursor clusters to more mature cells. When defining regulons that correlate with pseudotime, many of the regulons did not surpass a Pearson correlation coefficient of +/- 0.5, due to variable activity of regulons either within the middle of the trajectory or on either end (Supp Fig 3A). For example, KLF10 had high regulon activity in cluster C1 (central differentiation state) and decreased activity in both endpoints, to this end, correlation across the entire trajectory would not properly capture the changing activity across differentiation. To address this, we defined regulons changing across pseudotime for two trajectories (trajectory 1: C0,C1,C3,C4,C5 and trajectory 2: C1,C2,C6,C7) (Fig 3C). This analysis identified several regulons exhibiting a stronger correlation with pseudotime (Fig 3D, Supp Fig 3B-C).

For trajectory 1, the regulon of FOXP1, which regulates T cell quiescence and acts by inhibiting IL-7Rα, is strongly inversely correlated with pseudotime (r=-0.58, Pearson correlation, Supp Fig 3B, Fig 3D-E)^44^. Conversely, BCL3 was the most correlated regulon; as cells differentiate from the thymus, BCL3 regulon expression increases between clusters C0 and C1 and then decreases as cells differentiate along Trajectory 2 towards Th1/17/NK-like iNKTs (r=0.72, Pearson correlation, Fig 3D-E, Supp Fig 3B). Bcl3 is a pro-survival factor that contributes to immunologic tolerance and modulates NF-kB activity in mice^45,46^. Similarly, the regulon of ATF3, a negative regulator of IFNγ in murine NK cells^25^, is positively correlated with pseudotime along Trajectory 1 and then proceeds to decrease along trajectory 2. As ATF3 expression decreases from the Th2-like C1 to the more Th1-like C2/C6, we observe increased expression of IFNγ, providing a possible regulatory mechanism behind the increased activity of IFNγ in this iNKT population (Fig 3D, Fig 2E). Along trajectory 2, as cells differentiate between Th2 cells and Th1/17/NK-like iNKTs, the KLF10 regulon is dramatically reduced (KLF10 r= -0.61, Pearson), correlating with a role for KLF10 in restraining Th17-like differentiation described in another innate lymphocyte population^47^. The EOMES regulon increases along trajectory 2 (EOMES r= 0.68, Pearson), similar to the previously noted increase in EOMES between Th2-like and Th1/17/Nk-like iNKT cells in mice^18^. Taken together, these findings begin to elucidate the transcriptional regulatory networks that change as human iNKT cells differentiate.

### CD4 expression poorly delineates functional propensity in human iNKT cells

Due to the high dropout rate of *CD4* in scRNA-sequencing analyses, we utilized oligomer-conjugated antibodies to analyze CD4 and CD8 protein expression in addition to RNA expression (Fig 3F-G and Supp Fig 4A). While CD8^+^ human iNKT cells have been previously demonstrated by flow cytometry, this population is very small compared to CD4^+^ and double negative (DN, CD4^-^ CD8^-^) iNKT populations and little is known about its function^31^. Surprisingly, in this analysis, we found that CD8^+^ iNKT cells (C5) are characterized by a naïve/precursor-like transcriptional profile (Fig 3F-G). Most clusters expressed CD4, while part of C6 (C6.2) was DN (Fig 3F-G). While CD4 expression was originally attributed to iNKT2 cells in mice, subsequent reports indicated that CD4 expression could be present on a subset of Th1-like iNKT cells as well^11,48^. Similar to these murine reports, we found a CD4^+^ Th1/17/NK-like population in our human study (C2). This cluster had upregulation of *TBX21* and *RORC* along with many cytotoxicity and NK genes. These findings suggest that CD4 may not best distinguish functionally distinct subsets of iNKT cells.

**Figure 4.**
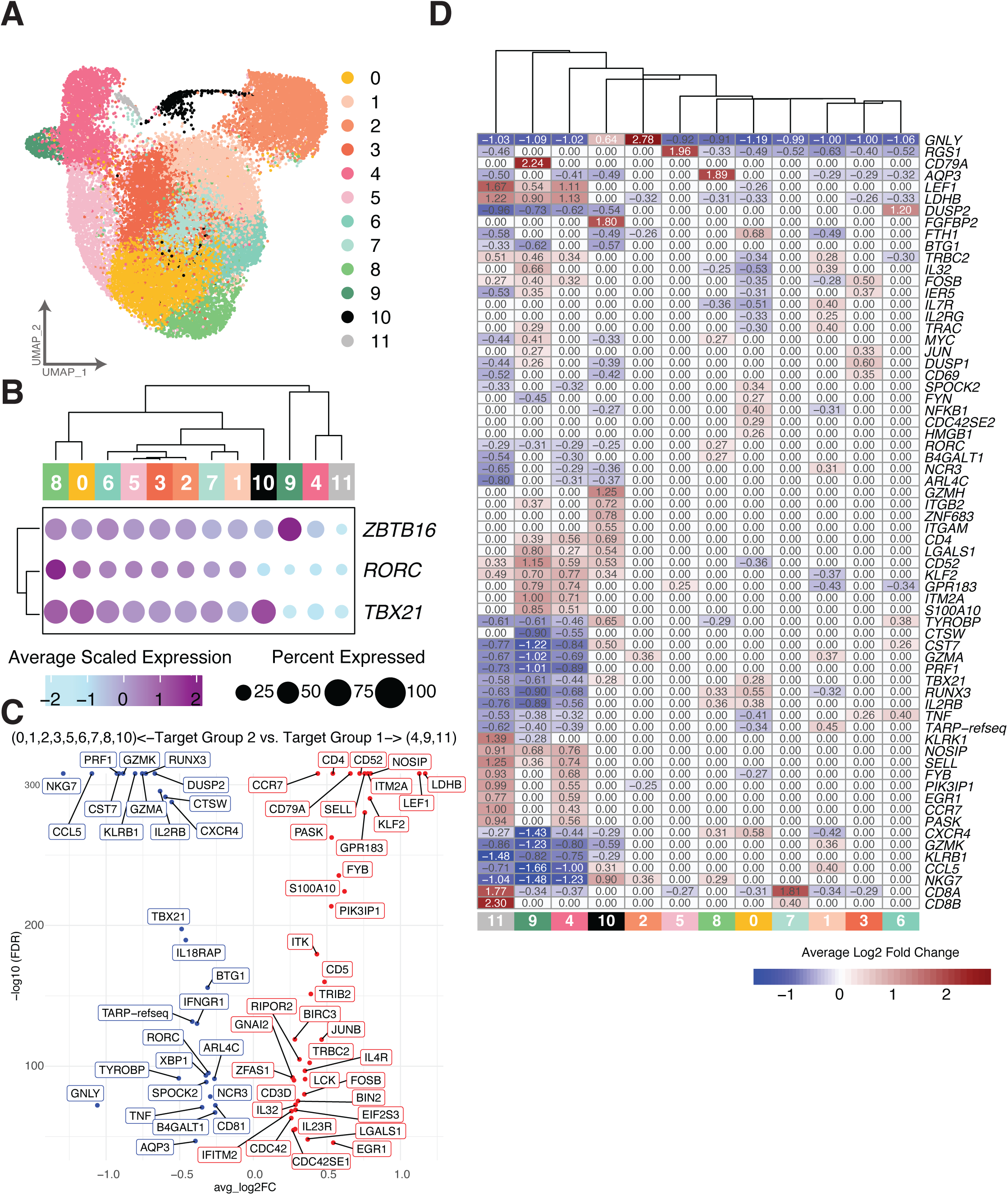
Transcriptional heterogeneity of human iNKT cells derived from peripheral blood. A) UMAP of all iNKT cells isolated from peripheral blood donors. Cells are colored by cluster assignment. B) Expression dot plots of key transcription factors. Size of dot indicates percent of cells expressing the indicated gene and color indicates the average scaled expression. C) Volcano plot of top differentially expressed genes separating PB-C4,9,11 from all other clusters. Each dot is labeled by the gene name and color of the dot and label indicates increased expression based on each of the target groups. D) Heatmap of the top 10 differentially expressed genes identified by each cluster. Each cell is colored by the average log2FC of each gene identified in each cluster. A value of 0 indicates the gene was not identified as being significantly up or down regulated for the labeled cluster.

### Transcriptional analysis of human peripheral blood-derived iNKT cells reveals further heterogeneity, particularly in the CD8^+^ population

Collectively, the WTA results on human iNKT cells derived from PB, CB, thymus, and BM suggest that subsets of human iNKT cells can be best described as Th1/17/NK-like, Th2-like, and naïve/precursor-like, based on expression patterns of transcription factors, naïve/memory phenotype, and genes important for cytotoxicity and NK cell function. We saw similar results in a transcriptional analysis employing a custom targeted panel of 435 genes (Supp Table 1). While we could not perform direct cell to cell comparison of the targeted assay to the WTA across the multi-tissue experiment, the overall correlation between the top variable genes in the WTA panel and targeted panel was very high (r=0.9, p<2.2e-16, Pearson correlation) and revealed a striking similarity in expression pattern of key genes despite having an overall smaller gene space (Supp Fig 5). To study PB-derived iNKT cell heterogeneity more robustly with higher cell numbers, we utilized this targeted scRNA-seq panel on 24,758 cells from 10 donors (Fig 4A, Supp Fig 6A).

**Figure 5.**
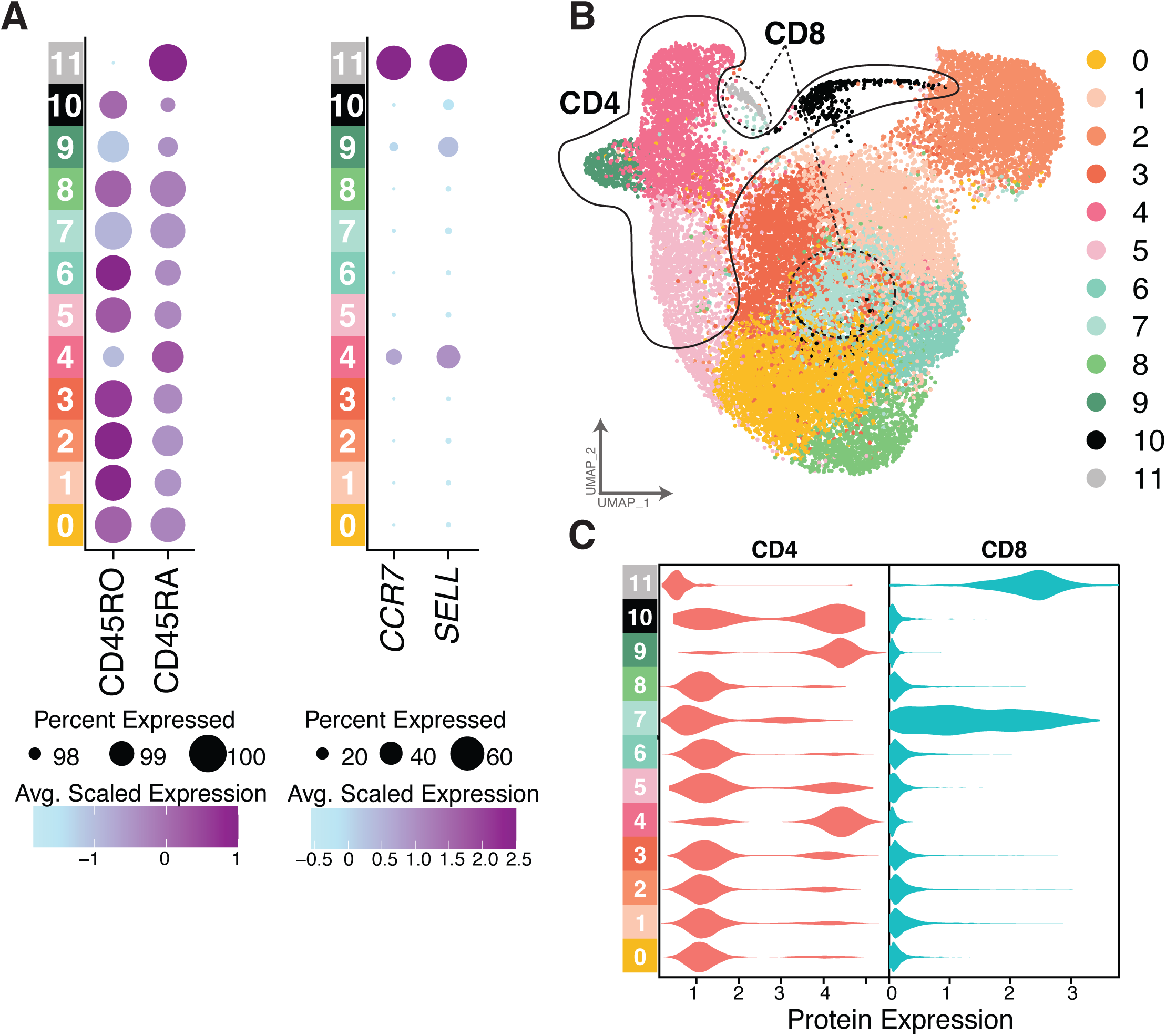
Naïve/memory and CD4/8 expression patterns of human peripheral blood iNKT cells facilitates detection of naïve/precursor-like and Th1/17/NK-like CD8^+^ iNKT cells. A) Dot plot of CD45RO and CD45RA protein expression and *CCR7* and *SELL* gene expression. For each figure, size of dot indicates percent of cells expressing the protein/gene of interest and color indicates average scaled expression. B) UMAP of iNKT cells colored by cluster assignment. Regions of UMAP are circled based on observed protein expression of CD4 and CD8. C) Violin plot of CD4 and CD8 protein expression separated by cluster.

**Figure 6.**
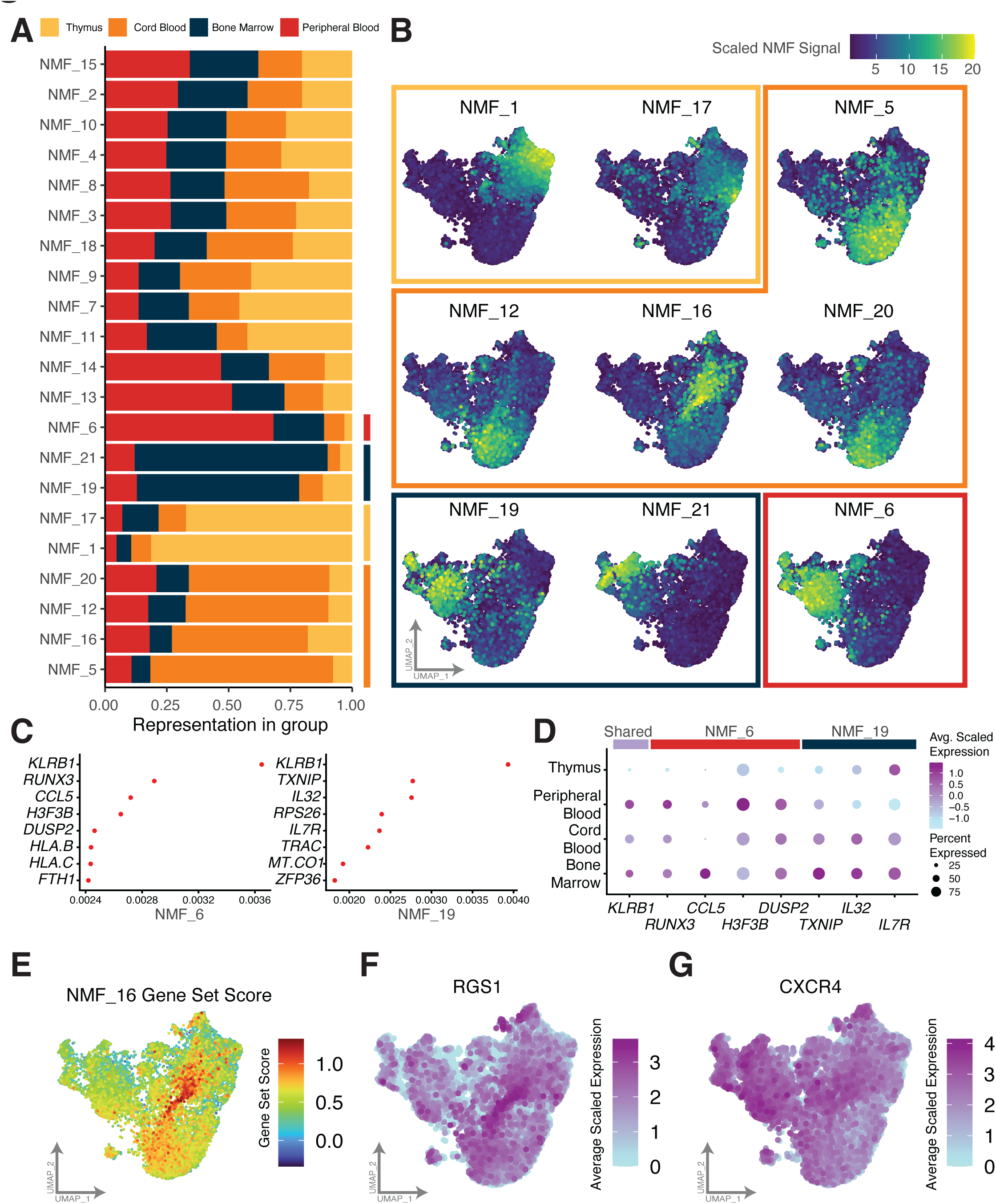
Transcriptional heterogeneity of human iNKT cells from different tissue sources within and between clusters. A) Barplot highlighting differing proportions of Non-negative matrix factorization (NMF) factors identified across each tissue type. The proportion of cells belonging to each factor (or biological process) is indicated by color. B) UMAP representation of iNKT cells colored by scaled NMF signal of 9 NMF factors that are highly represented by each tissue type. Each UMAP is grouped by square outline that is colored by the tissue type showing the strongest contribution. C) Visual loadings of the top eight key genes contributing to the two indicated NMF factors. D) RNA expression of a subset of genes identified as being key genes for two NMF Factors and their expression separated by tissue type. Size of dot indicates percent of cells expressing the indicated gene and color indicates the average scaled expression. E) Gene set expression of genes driving NMF_16 overlaying the iNKT UMAP. Each cell is colored by the scaled module score of the NMF_16 gene set. F-G) UMAP of iNKT cells colored by expression of annotated gene.

Similar to our previous analysis, we found naïve/precursor-like, Th2-like, and Th1/17/NK-like clusters within human PB iNKT cells (designated PB-C#) as based on transcription factor expression (Fig 4B); other differentially expressed genes including cytotoxicity genes and cytokine receptors (Fig 4C-D, Supp Fig 6B, Supp Table 5); and naïve/memory signatures (Fig 5A). Th1/17/NK-like clusters (PB-C0-3, PB-C5-8, and PB-C10) co-expressed *TBX21* and *RORC* (Fig 4B). These clusters also expressed the highest levels of genes important for cytotoxicity and NK function as shown in a volcano plot comparing them to the Th2-like cluster (PB-C9) and naïve/precursor-like clusters (PB-C4, PB-C11) (Fig 4C) and in a heatmap of the 10 most differentially expressed genes (Fig 4D). Naïve/precursor-like and Th2-like clusters were characterized by relative downregulation of these genes, with upregulation of genes characteristic of T cells, particularly Th2 cells, including *LEF1, ITK, CD52, CD4,* and *IL4R*. *ITK* has previously been shown to provide important signals for the differentiation of iNKT cells^49^. We again found that the Th2-like PB-C9 had high expression of *CD79A* (PB-C9: 2.24 avg log2FC), similar to our finding in the multi-tissue experiment (Fig 1F, 4D, Supp Table 5). Interestingly, we saw that the cluster with highest *RORC* expression (PB-C8) also had high and unique expression of *AQP3* (Fig 4D, 1.89 avg log2FC). *Aqp3* expression has also been demonstrated in iNKT17 cells in mice^17^.

We further utilized oligomer-conjugated antibodies to analyze CD4 and CD8 protein expression along with RNA expression. We found four clusters expressing CD4 (PB-C4-5 and PB-C9-10) (Fig 5B-C). PB-C9 also had the highest expression of *ZBTB16* (Fig 4B) and may represent Th2-like iNKT cells. Similar to previous reports and our prior analysis, we found two clusters (PB-C5 and PB-C10) among our Th1/17/NK-like clusters that were, to varying degrees, CD4^+^ (Fig 5C). While PB-C10 shares similar expression of many cytotoxicity and NK genes to the rest of the Th1/17/NK-like population, it also expressed *GZMB* and *GZMH* (Supp Fig 6B). This cluster was further characterized by the highest differential expression of *FGFBP2* (Fig 4D), the protein product of which has been previously reported to be secreted by cytotoxic T and NK cells^50^. Interestingly, we found two CD8^+^ iNKT clusters with very distinct transcriptional patterns—one within the naïve/precursor group of clusters (PB-C11), similar to our results in the multi-tissue experiment, and one in the Th1/17/NK-like group of clusters (PB-C7) (Fig 4B-D, 5A-B). Interestingly, PB-C11 has increased expression of both *CD8A* and *CD8B*, while PB-C7 has a much higher increase in *CD8A* expression relative to *CD8B* (Fig 4D, PB-C11 – *CD8A*: 1.77, *CD8B*: 2.30; PB-C7 – *CD8A*: 1.81, *CD8B*: 0.40 avg log2FC). These results suggest that at the protein level, iNKT cells in PB-C11 may express CD8αβ heterodimers, while iNKT cells in PB-C7 may express CD8αα homodimers, as has been demonstrated in NK cells with enhanced cytotoxicity and other innate lymphocyte populations^51–53^. Taken together, these data reveal heterogeneity within CD4^+^ and CD8^+^ human iNKT cell populations and suggest these markers may not be ideal correlates of iNKT cell function.

### Distinct biological processes differ by tissue type in iNKT cells

Although we found that PB-derived iNKT cells recapitulated the clusters observed in the multi-tissue experiment, we sought to further evaluate whether differences between iNKT cell populations from each tissue type could be detected, possibly even within the same clusters. We performed Non-negative Matrix Factorization (NMF) to evaluate biological processes that are differentially represented in each tissue type. 21 different factors (or biological processes) were identified with factors NMF_1 and NMF_17 being highly represented in thymic cells deriving predominantly from C0 and C5 (Fig 6A-B). CB processes included NMF_5, NMF_12, NMF_16 and NMF_20 spanning C0 and C1. NMF_19 and NMF_21 are enriched in BM cells from C2 and C6 while NMF_6 is over represented in PB from the same two clusters. All other gene programs were relatively evenly distributed among cells derived from each tissue type.

NMF_6 (PB) and NMF_19 (BM) occupy the same clusters (C2 and C6) yet contain different genes driving each biological process. While *KLRB1* is the primary gene associated with both factors, NMF_6 is strongly driven by *RUNX3, CCL5, H3F3B* and *DUSP2* while NMF_19 is associated with *TXNIP, IL32* and *IL7R* (Fig 6C-D). RNA expression across all iNKT cells reveals that the top genes for each process are largely associated with either PB or BM. These findings suggest that while most features of the Th1/17/NK-like clusters are largely shared by the PB and BM derived iNKT cells, they contain underlying biological gene programs that distinguish them.

With respect to the thymus and CB, the biological process NMF_16 overlays a connecting population of cells that very clearly overlaps the Monocle pseudotime trajectory between the naïve/undifferentiated C0 and Th2-like C1 clusters (Fig 6B, Fig 3C). C0 and C1 are largely populated by cells derived from the thymus and CB (Fig 1E). Genes driving this biological process include *RGS1, CXCR4, CD69, DDX21, BTG1, JUN* and *KLF6*, among 19 others. When evaluating the joint expression of all 19 genes driving this biological process using a gene set score, we observe that cells spanning the junction between C0 and C1 are clearly highlighted (Fig 6E). Evaluating specific genes separately, *RGS1* is highly expressed at the same junction while *CXCR4* is more globally expressed across various cells, albeit still showing high expression at the C0-C1 junction (Fig 6F-G). Using the ingenuity pathway analysis to evaluate genes in NMF_16, we found that both *JUN* and *CXCR4* are mapped to the CXCR4 signaling pathway. The CXCR4-CXCL12 signaling axis is critical for cell development and migration in several lymphocyte populations in both humans and mice. Mice with CXCR4 deficiencies had a reduced abundance of functional NK cells in various tissues including peripheral blood, bone marrow and spleen^54^. Further *Rgs1* expression in mouse thymic iNKT cells was briefly noted to be upregulated during cell development and maturation^55^. *Klf6* expression has been shown to be increased upon acquisition of a memory phenotype in CD4^+^ T cells^56^. Taken together, this biological process is likely describing the early transcriptional changes that are driving iNKT cells’ maturation and egress from thymus. By integrating unstimulated iNKT cells across tissue types and evaluating shared regulatory processes these data begin to piece together the underlying transcriptional mechanisms driving iNKT cell development and heterogeneity.

## DISCUSSION

iNKT cells represent an evolutionarily conserved critical component of immune responses in a variety of disease states as well as a novel approach to universal donor cellular therapies. However, a better understanding of human iNKT cell biology is required to realize the full potential of these cells. Herein, we report an integrated transcriptional analysis of human iNKT cells from peripheral blood, cord blood, thymus, and bone marrow, and reveal critical insights into the heterogeneity of these cells in these immunologically relevant tissues.

Our data demonstrate that human iNKT cells form a differentiation trajectory which, when limited to post-stage 0 iNKT cells, appears to originate from a naïve/precursor stage. The reduced expression of *ZBTB16* in our naïve/precursor stage relative to the Th2-like clusters corresponds with a recent report of a similar population in murine thymic iNKT cells^17^. Further, our naïve precursor stage was characterized by expression of genes known to play important roles in murine iNKT cell development. *Lef1* is expressed in ST0 and early differentiating iNKT cells and has also been implicated in iNKT2 differentiation in mice^16,20,38^. *Sox4* is also important for iNKT cell development by inducing microRNA-181 to enhance TCR signaling in iNKT precursors^39^. *Satb1* appears to regulate lineage defining factors in certain post-selection thymocytes, including iNKT cells^40^. These cells were also characterized by increased *CCR7* expression, which has been previously associated with a more immature phenotype of iNKT cells in mice^57^. While the expression patterns of CD45RA and CD45RO in human iNKT cell subsets remain poorly understood, the use of oligomer conjugated antibodies allowed us to determine that expression of CD45RA in these *CCR7*-expressing precursor clusters corresponds with the canonical definition of human naïve conventional T cells (CD45RA^+^CCR7^+^). Most of the remaining iNKT cells in our study express a T effector memory profile (CD45RA^-^*CCR7*^-^). However, we also found a small group of DN iNKT cells displaying a TEMRA-like expression pattern (CD45RA^+^*CCR7*^-^). Among conventional T cells, TEMRA cells are mostly CD8^+^, with the highest IFNγ production and cytotoxic function, and increase in PB and BM with age^58–60^. Rare CD4^+^ TEMRA cells also demonstrate cytotoxic function^61^. Thus, TEMRA iNKT cells share many similarities with TEMRA conventional T cells.

Previous work has suggested that murine iNKT cells emerge from the thymus at multiple points along the differentiation trajectory^16,62^. Accordingly, we found that all naïve/precursor clusters were populated by PB iNKT cells in addition to thymic iNKT cells. We further found that Th2-like iNKT clusters were predominated by CB iNKT cells, yet contained PB iNKT cells as well. This corresponds with previous data showing that human CB derived iNKT cells produce higher Th2 cytokines than iNKT cells derived from adult PB^34,63^. Our differentiation trajectory suggests that Th2-like iNKT cells represent a transitional phenotype between naïve/precursor iNKT cells and Th1-like iNKT cells, similar to recent findings in murine iNKT cells^16,17^. In these murine studies, iNKT2 cells (or a fraction thereof) represented a transitional population from which iNKT1 cells and iNKT17 cells independently branched off. Indeed, several murine studies have highlighted the distinct transcriptional clustering of iNKT1 cells and iNKT17 cells, with unique expression of *Tbx21* and *Rorc*, respectively, which has been further demonstrated at the protein level^12,16,17,20^. In contrast, our study of human iNKT cells revealed co-expression of *TBX21* and *RORC* in two clusters, suggesting a combined Th1/17 signature in these cells.

Our analysis has started to tease out the complex network of transcription factors that work in concert to alter the differentiation state of human iNKT cells. We highlighted several novel transcription factors that change over the course of differentiation including *BCL3*, *FOSL2*, and *FOXP1*. Notably, B cell lymphoma 3 protein (BCL3) modulates cell-cell interactions in the stroma supporting immunologic tolerance^45,46^. Understanding if these regulons and transcription factors are altered under different conditions and exploring cell-cell interactions in a single cell manner will start to highlight how mechanistically the regulatory network is altering the cellular environment. Furthermore, we observed that FOXP1 regulon activity was inversely correlated with early differentiation of iNKT cells. FOXP1 competes with FOXO1 for *Il7r* enhancer binding and leads to transcriptional repression, suggesting that de-repression of *Il7r/IL7R* may be required for iNKT maturation^44^.

The most differentiated cells in our analysis, with the highest expression of genes important for cytotoxicity and other effector functions, were predominantly BM and PB derived. As our tissues of origin also reflect distinct timepoints across the age spectrum (with CB representing neonates, thymus acquired from children, and PB and BM specimens from adult donors), we cannot exclude an age-dependent influence on our findings. Indeed, multiple murine and human studies have suggested changes to the iNKT cell population with aging, with a progression from a Th2-like iNKT cell predominance early in life to greater relative abundance of Th1-like iNKT cells over time^33,34,63,64^. However, it remains important to note that all described clusters were found in adult PB samples. The Th1/17/NK-like population was transcriptionally characterized by many features described in other cytotoxic innate lymphocytes. This included expression of receptors for cytokines known to be expressed by NK cells^32^. In addition, the expression of granzyme K was recently reported to be a hallmark of innate lymphocytes, and interestingly activated by cytokine stimulation rather than TCR activation^26^.

While many studies of iNKT cells compare CD4^+^ to CD4^-^CD8^-^ (DN) cells, CD8^+^ iNKT cells are a relatively understudied population, likely owing to their rarity in humans and absence in mice^31,65,66^, though this absence in mice has been called into question^67^. As such, the developmental program of CD8^+^ iNKT cells remains unclear, and whether CD8^+^ iNKT cells arise directly from double positive Stage 0 iNKT cells through loss of CD4 or whether CD8 is re-expressed at some point in the developmental trajectory remains unknown. Our data showing a CD8^+^ cluster with a naïve/precursor profile suggests the former, though further study is required to definitively answer this question. We detected a second CD8^+^ cluster with a Th1/17/NK-like transcriptional profile in our PB only experiment, but not in our multi-tissue experiment, which may be related to substantially different numbers of iNKT cells analyzed between the two experiments. Similar to previous murine findings that iNKT1 cells can be CD4^+^ or CD4^-11,48^, we found CD4^+^ cells within the Th1/17/NK-like clusters. Our data suggest that the presence or absence of CD4 and CD8 expression does not clearly delineate functionally distinct subsets.

Altogether our results reveal critical insights into the transcriptional heterogeneity of human iNKT cells. Defining the transcriptional profile of human iNKT cells in homeostasis will facilitate future experiments to understand changes to these transcriptionally distinct subsets in various disease states or during *ex vivo* expansion. Future work will also be required to determine in what immune contexts they are differentially activated, which may prove tissue specific. Further, our analysis of unstimulated cells will support the future development of an iNKT cellular therapy platform using iNKT cell subsets with distinct functional profiles. Although future studies are required to translate our transcriptional findings into a demonstration of differential function, and additional modalities such as assay for transposase-accessible chromatin with sequencing (ATAC-seq) could further refine our understanding of the regulatory network underlying iNKT cell differentiation, these data lay the groundwork for the development of optimized iNKT-based cellular therapies. Novel therapies harnessing the universal donor potential of these innate lymphocytes may be enhanced in safety and efficacy by targeting the subset displaying the desired function. Herein we provide a critical first step towards realizing that translational goal.

## MATERIALS AND METHODS

### Tissue Acquisition and Preparation of Single Cell Suspensions

This study was determined by the Stanford University Institutional Review Board to be IRB-exempt as secondary research on de-identified samples acquired through IRB-approved research protocols. Thymic tissue was acquired from congenital cardiac disease surgery patients (excluding those with Chromosome 21q11.2 Deletion (DiGeorge) syndrome or other immune disorders) and was mechanically dissociated prior to downstream processing. Bone marrow was acquired from healthy bone marrow transplant donors, resuspended to dissociate clumps, and filtered to remove bone fragments. Peripheral blood was acquired from healthy adults donating platelets (leukoreduction filter chambers). Cord blood was collected from donors at the time of parturition. All specimens underwent Ficoll-Paque density gradient centrifugation and were then cryopreserved until ready for use. Peripheral blood samples for the peripheral blood-only experiment were acquired on the same day and not frozen.

### Human iNKT cell enrichment and purification

Human iNKT cells were enriched by magnetic bead-based positive selection (Stem Cell Technologies EasySep™ Release Human PE Positive Selection Kit) utilizing PE-conjugated anti-Vβ11 antibody (Beckman Coulter, C21), followed by bead release, additional staining with fluorochrome-conjugated and oligomer-conjugated antibodies, and fluorescence activated cell sorting for CD24^-^CD3^+^Vβ11^+^Vα24^+^ cells (4-way purity setting on FACSAria II). Antibodies included APC-H7 CD14 (BD Biosciences, MoP9), APC-Cy7 CD19 (BD Biosciences, SJ25C1), APC-Cy7-CD24 (Biolegend, ML5), PerCP-Cy5.5 CD3 (Biolegend, UCHT1), FITC Vα24 (Beckman Coulter, C15), PE Vβ11 (Beckman Coulter, C21), Oligo-CD4 (BD AbSeq, SK3), Oligo-CD8 (BD AbSeq, RPAT8), Oligo-CD45RA (BD AbSeq, HI100), and Oligo-CD45RO (BD AbSeq, UCHL1). Samples were sorted individually, cell counts roughly normalized, and then pooled for sequencing. scRNA-seq libraries were generated using the Rhapsody platform (Becton Dickenson) employing a targeted panel (PB only) or targeted panel and whole transcriptome analysis (multi-tissue experiment). Data was processed using the BD Rhapsody pipeline (Seven Bridges) followed by analysis in R.

### scRNA-seq Data Preprocessing for Multi-Tissue Whole Transcriptome Data

Expression tables (RSEC_MolsPerCell.csv) and sample tags (Sample_Tag_Calls.csv) were downloaded from Seven Bridges and analyzed using Seurat (v4.4.0). First, separate assays were created using the CreateSeuratObject function and percent mitochondrial DNA, nCount_RNA and nFeature_RNA were evaluated for each cell. Cells were maintained for downstream processing if they met the following criteria: nFeature_RNA > 200; nFeature_RNA < 4000; nCount_RNA < 20000; percent.mito < 20. Cells with sample tags labeled “Undetermined” or “Multiplets” were filtered out. Each sample was scaled and normalized using Seurat’s ‘SCTransform’ function to correct for batch effects (with parameters: vars.to.regress = c("nCount_RNA"), dims = 1:20). Any merged analysis or subsequent subsetting of cells/samples underwent the same scaling and normalization method. Cells were clustered using the original Louvain algorithm and top 10 PCA dimensions via ‘FindNeighbors’ and ‘FindClusters’ (with parameters: resolution = 0.5) functions. After initial subsetting and clustering, a clear population of infiltrating macrophages were identified exhibiting higher expression of *FCGR3A, LYN, FCR1G, IL1B*. The macrophage cluster was filtered out and we subsequently used Harmony (group.by.vars = "Sample_Tag", reduction = "pca", assay.use = "SCT") before rerunning RunUMAP, FindNeighbors and FindClusters to annotate final object (dims=1:20). The resulting merged and normalized matrix was used for the subsequent analysis. To evaluate cluster resolution we utilized clustree before settling on cluster assignment. ADT data was normalized using NormalizeData command with method = “CLR” and a margin of 2.

### scRNA-seq Data Preprocessing for Multi-Tissue and Peripheral Blood-only Targeted Panel Data

Expression tables (MolsPerCell.csv) and sample tags (Sample_Tag_Calls.csv) were downloaded from Seven Bridges and analyzed using Seurat (v4.4.0). RSEC_MolsPerCell file was used for the Multi-tissue targeted panel and DBEC_MolsPerCell used for the peripheral blood-only targeted panel. First, separate assays were created using the CreateSeuratObject function. Each sample was scaled and normalized using Seurat’s ‘SCTransform’ function to correct for batch effects (with parameters: vars.to.regress = c("nCount_RNA"), dims = 1:20). Any merged analysis or subsequent subsetting of cells/samples underwent the same scaling and normalization method. Cells were clustered using the original Louvain algorithm and top 10 PCA dimensions via ‘FindNeighbors’ and ‘FindClusters’ (with parameters: resolution = 0.5) functions. After initial subsetting and clustering, a clear population of infiltrating macrophages were identified exhibiting higher expression of *FCGR3A, LYN, FCR1G, IL1B*. The macrophage cluster was filtered out along with cells with sample tags labeled “Undetermined” or “Multiplets”. Only the Multi-Tissue sample subsequently used Harmony (group.by.vars = "Sample_Tag", reduction = "pca", assay.use = "SCT"), while both datasets underwent the same downstream preprocessing using RunUMAP, FindNeighbors and FindClusters to annotate final object (dims=1:20 for multi tissue data and dims=1:10 for peripheral blood data). The resulting merged and normalized matrix was used for the subsequent analysis. To evaluate cluster resolution we utilized clustree before settling on cluster assignment. ADT data was normalized using NormalizeData command with method = “CLR” and a margin of 2.

### Differential scRNA Expression Analyses

For cell-level and cluster-level differential expression, we used the ‘FindMarkers’ or "FindAllMarkers’ Seurat function as appropriate, with a minimum pct. of 0.10. The resulting differentially expressed genes (DEGs) were then filtered for adjusted p-value < 0.05 and sorted by fold change. All differential expression analyses were carried out using the "SCT" assay.

### Monocle pseudo-time analysis

Cell state decisions and trajectory-based analyses were constructed by Monocle3. All cells from different samples were extracted and imported into Monocle3 for analysis. Parameters for the analysis are consistent with the tutorial by the tool’s developers (http://cole-trapnell-lab.github.io/monocle-release/docs/#constructing-single-cell-trajectories). For the order_cells function, the thymic high cluster was used as the root node. For the learn_graph function ncenter=850 was used to learn the final trajectory. Monocle pseudotime scores were carried over to the original Seurat object for differential gene expression analysis.

### CytoTRACE

CytoTRACE was run using the R implementation (https://cytotrace.stanford.edu/) on the RNA counts matrix (v0.3.3). Resulting values were added to metadata for plotting using Seurat.

### SCENIC Regulon Analysis

RNA count matrices were extracted from the subsetted seurat object and run using the parameters of the Docker pySCENIC implementation v0.12.1 (https://pyscenic.readthedocs.io/en/latest/installation.html). Mitochondrial annotated genes and genes with low overall cell counts (<=1% of cells expressing the gene) were filtered prior to running pySCENIC. pySCENIC was run using 11 iterations and AUC scores were recalculated for regulons present in at least 60% of runs. The final SCENIC activity scores were added to the Seurat object as a new assay and scaled before evaluating using FindAllMarkers for differentially expressed regulon activity (logfc.threshold = 0.005).

### Ingenuity Pathway Analysis (IPA)

For all differentially expressed genes (Supp Table 2) we utilized ingenuity pathway analysis core analysis module to identify pathway enrichment in each cluster. Full results of the pathway analysis are available in Supplementary Table 4.

### Non-negative Matrix Factorization (NMF)

To run NMF we used the R package singlet (v.0.0.99, https://github.com/zdebruine/singlet)^68^. NMF gene programs were visualized using the UMAP reduction derived from SCT gene expression data.

### Add Module Score

We first identified the top genes driving NMF_16 and then used the AddModuleScore function in Seurat using the following gene set: "AL390957.1","SLC2A3","PRRC2C","CYTIP","JMJD1C","ELF1","SMAP2","YPEL5","FYB1","SOX4"," PTGER4","NDFIP1","IRF2BP2","STK4","VAMP2","IL6ST","TSC22D3","STK17B","ZBTB10","JUN","T NFAIP3","ITM2A","KLF6","SARAF","BTG1","DDX21","WHAMM","CD69","CXCR4","RGS1".

## ACKNOWLEDGEMENTS

We wish to thank the Stanford University Shared FACS Facility; sorting was performed on instruments in the core facility, including those obtained using an NIH S10 Shared Instrument Grant (S10RR025518-01). We also wish to thank Bita Sahaf, PhD, the Stanford Human Immune Monitoring Center, and the Texas A&M AgriLife Research Genomics & Bioinformatics Service.

R.G.J. received funding from the Washington University SPORE in Pancreatic Cancer Grant (P50CA272213). C.B. was supported by a Postdoctoral Research Training T32 Fellowship with support from Ruth L. Kirschstein Institutional National Research Service Award. This work was supported by the National Institutes of Health: P01 HL075462 to R.S.N and K08 HL151809 to M.M. M.M. also received funding for this work from the St. Baldrick’s Foundation (Scholar Award with generous support from the Rays of Hope Hero Fund).

## SUPPLEMENTARY MATERIALS

**Supplementary Figure 1.** A) UMAP representation of iNKT cells derived from multi-tissue samples. Each UMAP is separated by donor and colored by cluster assignment.

**Supplementary Figure 2.** A) Selected pathways identified by Ingenuity Pathway Analysis as being differentially upregulated in each cluster. Each pathway shows the relative -log(p-value) on the axis and the color of each bar indicates the z score. Positive z-score values indicate activation, while negative values indicate inhibition. B) Expression plots of three subsets of genes. Size of dot indicates percent of cells expressing each gene and color indicates the average scaled expression.

**Supplementary Figure 3.** A) Regulons correlating with pseudotime across all cells. B) Regulons correlating with pseudotime across cells within trajectory 1. C) Regulons correlating with pseudotime across cells in trajectory 2.

**Supplementary Figure 4.** A) Feature Plots indicating expression of the listed genes. Each cell is colored by the average scaled expression.

**Supplementary Figure 5.** A) UMAP of targeted data from the multi-tissue dataset. Each cell is colored by cluster assignment. B) Correlation of the average expression of 382 variable genes overlapping both the whole-transcriptome data and targeted data. Pearson correlation value and p value are indicated on the plot.

**Supplementary Figure 6.** A) UMAP representation of iNKT cells derived from peripheral blood samples. Each UMAP is separated by donor and colored by cluster assignment. B) Expression plots of a subset of genes. Size of dot indicates percent of cells expressing each gene and color indicates the average scaled expression.

**Supplementary Table 1.** Targeted scRNA-seq panel genes

**Supplementary Table 2.** Differentially expressed genes identified by cluster in the whole-transcriptome multi-tissue iNKT cell analysis.

**Supplementary Table 3.** Differentially expressed regulons identified by cluster in the whole-transcriptome multi-tissue iNKT cell analysis.

**Supplementary Table 4.** Ingenuity Pathway Analysis results of differentially expressed genes for WTA multi-tissue data.

**Supplementary Table 5.** Differentially expressed genes identified by cluster in the targeted peripheral blood iNKT cell analysis.

